# Category-induced global effects of feature-based attention in human visual system

**DOI:** 10.1101/2022.12.21.521513

**Authors:** Ling Huang, Jingyi Wang, Qionghua He, Chu Li, Yueling Sun, Carol A. Seger, Xilin Zhang

**Author notes:** These authors contributed equally to this work (Ling Huang, Jingyi Wang). Correspondence, Xilin Zhang, Professor, Key Laboratory of Brain, Cognition and Education Sciences, Ministry of Education, School of Psychology, South China Normal University, Guangzhou, Guangdong, China, Website: https://xlzhanglab.scnu.edu.cn/.

## Abstract

Global effects of FBA are generally limited to stimuli sharing the same or similar features, as hypothesized in the “feature-similarity gain model”. Visual perception, however, often reflects categories acquired via experience; whether the global-FBA effect can be induced by the categorized features remains unclear. Here human subjects were trained to classify motion-directions into two discrete categories and performed a classical motion-based attention task. We found a category-induced global-FBA effect in both the MT+ and frontoparietal areas, where attention to a motion-direction globally spread to unattended motion-directions within the same category, but not to those in a different category. Effective connectivity analysis showed that the category-induced global-FBA effect in MT+ was derived by feedback from the IFJ. Altogether, our results reveal for the first time a category-induced global-FBA effect and identify a source for this effect in human prefrontal cortex, implying that FBA is of greater ecological significance than previously thought.

---

Although a substantial amount of research suggests that a primary unit of attentional selection is spatial location, increasing evidence has demonstrated that attention can also select specific features independent of their spatial locations, known as feature-based attention (FBA)^1,2^. FBA not only modulates the neural response to different feature dimensions in their specialized cortical modules^3–10^ but can also select different feature values within a particular dimension, such as an orientation^11–15^, a color^16,17^, and a direction of motion^18–22^. Previous studies have indicated that FBA plays a key role in identifying and highlighting a target in a complex scene since we often have a priori knowledge of a target-defining feature but not of its exact location. The capacity of FBA to quickly select a target is mainly attributed to its global effect, as proposed by the “feature-similarity gain model”^21^, whereby FBA modulates the gain of cortical neurons tuned to the attended feature and those in its close vicinity, not only at an attended location but throughout the whole visual field.

The global nature of FBA has been supported by numerous psychophysical^13,16,23–25^, neurophysiological^19,22,26–28^, electroencephalogram (EEG)^17,29,30^, magnetoencephalogram (MEG)^11,31^, and functional magnetic resonance imaging (fMRI)^12,20,32,33^ studies using a variety of different types of stimuli. These studies have generally limited their scope to examining global FBA effects resulting from stimuli that share the same or highly similar features with the attended stimulus. However, visual perception is often categorical, reflecting various learned categories of objects (e.g., food, animals, plants) ^34–43^. These categories are critical for our brain in assigning ecological meaning and selecting adaptive behavioral responses to a stimulus^44,45^. Therefore, we proposed that relative to feature similarity, it would be advantageous for global FBA to extend to stimuli in the same category, particularly in cluttered scenes.

How does a categorized feature induce a global FBA effect in the visual system? We hypothesized that this effect would result from top-down feedback from frontoparietal cortical areas to visual areas. Several neurophysiological studies have identified some neurons in frontoparietal cortical areas that have very large receptive fields^26,28^. Although the receptive fields of these neurons are not centered in the ipsilateral hemifield, many do extend into the ipsilateral hemifield, especially with longer stimulus presentation times^46,47^. These large receptive fields may provide a critical underlying neural basis that can support a category-induced global effect of FBA. Consistent with this theory, a number of previous studies have revealed that the frontoparietal network plays a key role in the feature similarity-based global effect of FBA^10,26,28,33,48^. Intriguingly, the same frontoparietal networks have been shown to underlie the representation of category membership in category learning, as indicated by previous neurophysiological^36,43,49–54^ and brain imaging^55–58^ studies. This common role of the frontoparietal network in both global FBA and categorization supports the hypothesis that categorical representations could be a source of global FBA effects. Attending to a feature from a category would elevate the neural response to this category in frontoparietal cortical areas and result in active and global top-down biasing signals to the representations of other features falling within the same category.

To test this hypothesis, we trained human subjects to classify motion-directions into two discrete categories (Fig. 1b). Before and after this training, subjects were asked to perform a classical motion-based attention task and we measured both the motion aftereffect (MAE, Fig. 1d) and blood oxygenation level-dependent (BOLD) signals (Fig. 1e) evoked by the unattended motion-direction, either in the same or different categories as the attended motion-direction. Mechanistically, we performed interregional correlation and effective connectivity analyses to identify the neural mechanism underlying the category-induced global FBA effect.

**Fig. 1.**
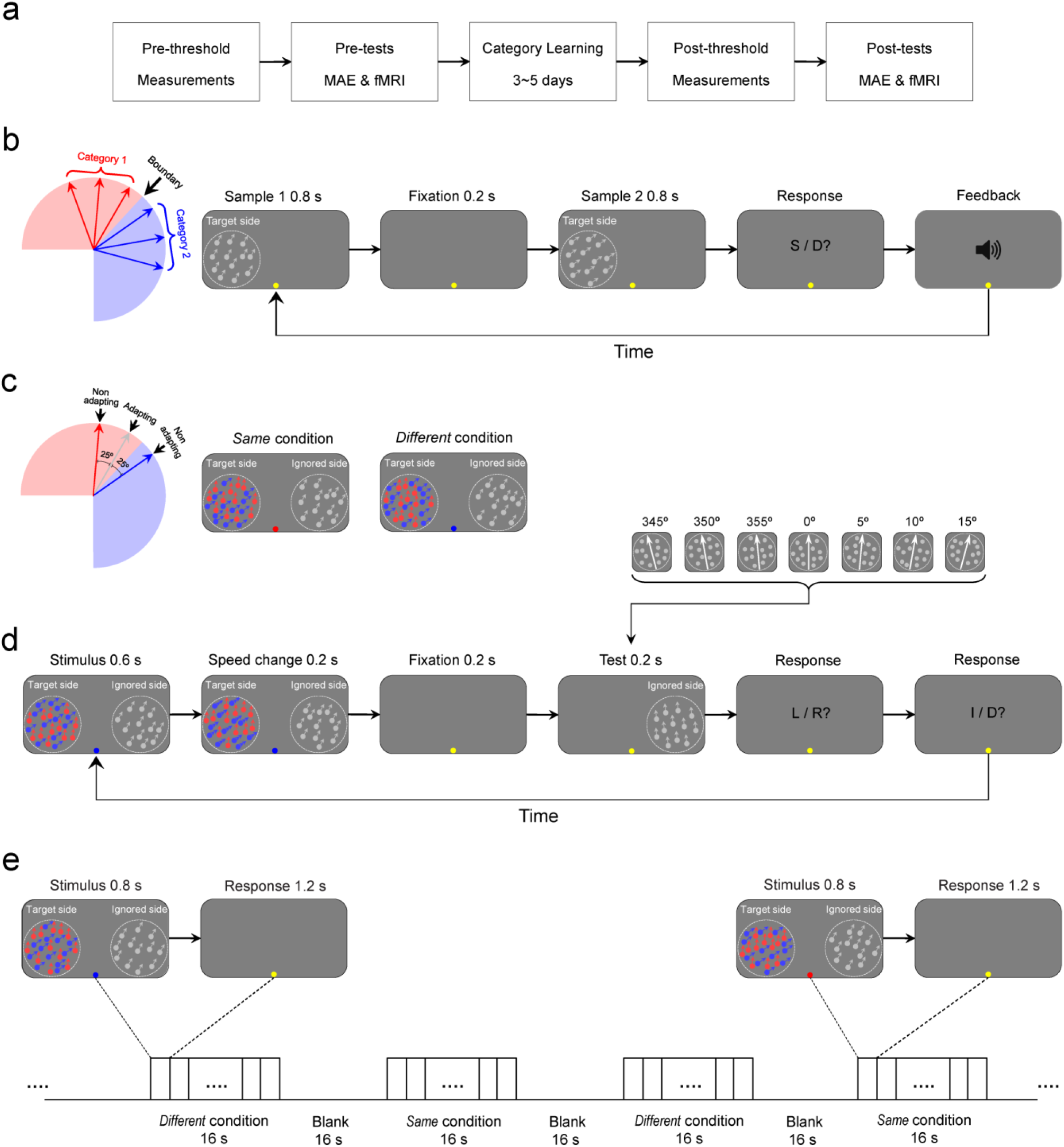
Stimuli and procedures in psychophysical and fMRI experiments. **a** Order of the procedures across the entire experiment. Note that the pre-tests included two separate MAE sessions for the *Same* and *Different* conditions and two fMRI sessions (retinotopic mapping and the FBA pre-test). The post-tests repeated all of these measurements with the exception of the retinotopic mapping fMRI session. **b** Motion-directions, categories, and the procedure for the category learning task. Left: the black arrow points to the motion-direction of the category boundary (in this case, the category boundary was 42.5°), which was randomly selected from 17.5°, 42.5°, 317.5°, and 342.5° clockwise from the vertical for each subject. The motion-directions in the red zone belonged to one category, and those in the blue zone belonged to another category. Right: procedure for each trial of the category learning task. **c** Three sample motion-directions used in the threshold measurements and motion aftereffect tasks: the adapting motion-direction (gray arrow), a non-adapting motion-direction belonging to the same category as the adapting motion-direction (*Same* category condition, red arrow), and the other non-adapting motion-direction belonging to the other category (*Different* category condition, blue arrow). Note that the colored dots here are for illustration purposes only; they were not used in the experiments. The angular difference (25°) between the adapting motion-direction and the same category motion-direction was equal to that between the adapting motion-direction and the different category motion-direction. The colored fixation indicate the attended motion-directions (the attended stimulus) on the target side: red and blue indicated the *Same* and *Different* category conditions, respectively. **d** Stimuli and procedure for the motion aftereffect (MAE) task. On each trial, the stimulus was presented for 0.6 s, followed by a 0.2-s speed change of the attended stimulus on the target side and a 0.2-s fixation interval. Then one of the 7 test stimuli (0°, 5°, 10°, 15°, 345°, 350°, and 355° clockwise from the vertical) was randomly presented on the ignored side for 0.2 s, followed by two response intervals. Subjects were first asked to make a 2AFC judgment on the motion direction of the test stimulus on the ignored side, either leftward or rightward. Then, subjects needed to make another 2AFC judgment on the speed change of the attended stimulus on the target side, either increased or decreased. Double arrows indicate the increased speed of the dots. L/R: leftward or rightward; I/D: increased or decreased. **e** Stimuli and procedure used in the fMRI experiments. Subjects attended one direction of motion (the attended stimulus) within a display of overlapping dots on the target side, and ignored the unattended stimulus on the ignored side. The attended stimulus on the target side was indicated by a colored fixation dot: red and blue indicated the *Same* and *Different* conditions, respectively. On each trial, the stimulus was presented for 0.8 s, followed by a fixed 1.2-s fixation interval, and subjects did a 0.2-s speeded discrimination at threshold.

## Results

### Experiment overview

In this study, we examined whether a global FBA effect can be induced by a learned categorical structure of stimulus features. The experiment consisted of 5 stages conducted across multiple days: threshold pre-measurements ➔ MAE & fMRI pre-tests ➔ category learning (3~5 days) ➔ threshold post-measurements ➔ MAE & fMRI post-tests (Fig. 1a), and each stage consisted of multiple sessions. Before the category learning task, subjects completed separate sessions on different days for (1) threshold testing in the *Same* condition (2) threshold testing in the *Different* condition (3) the MAE task in the *Same* condition (4) the MAE task in the *Same* condition (5) fMRI retinotopic mapping scans and (6) fMRI FBA scans. The order of these sessions varied, but threshold testing sessions were always first (with order counterbalanced across subjects), and the order of the two MAE sessions was counterbalanced across subjects. Subjects then completed intensive category learning which continued until criterion performance was reached (3-5 days of training). After the category learning task, subjects repeated all the same tasks as in the pretraining sessions (threshold testing, MAE tests in the *Same* and *Different* conditions, and fMRI FBA scanning session), with the exception of the retinotopic mapping scan which was performed only during pretraining.

### Category learning

In the category learning stage, we trained human subjects to classify motion directions into two discrete categories on the target side. For each subject, the category boundary motion-direction was randomly selected from 17.5°, 42.5°, 317.5°, and 342.5° clockwise from the vertical. Two groups of 3 motion-directions were separated by a category boundary that centered the range of the motion-directions. Motion-directions were evenly spaced (25° apart, Fig. 1b). For each trial, the first sample stimulus (in which the dots moved in 1 of 6 evenly spaced directions) was presented for 800 ms, followed by a 200 ms fixation interval. Then, the second sample stimulus (in which the dots moved in 1 of the remaining non-identical 5 motion-directions) was presented for 800 ms and was followed by a response interval. Subjects were asked to press one of two buttons to indicate whether the second sample stimulus was in the same or different category as the first sample stimulus and received auditory feedback if their response was incorrect. The training was terminated when the subject’s accuracy met a criterion of greater than 85% for each of the 6 motion-directions. The criterion was assessed at the end of each 120-trials block, and was applied to each of the 6 motion-directions individually (about 18.0 ± 1.31 blocks).

### Psychophysical experiments: motion aftereffects

Before and after category training, we measured the MAE resulting from adaptation to an unattended motion-direction on the ignored side during separate *Same* and *Different* conditions. In the *Same* condition, the category of the attended stimulus on the target side matched the category of the unattended stimulus on the ignored side; in the *Different* condition the categories did not match. The MAE task was performed in two separate sessions (the *Same* and *Different* conditions) occurring on different days; the order of the two sessions was counterbalanced across subjects. During each session, the attended stimulus on the target side was indicated by a colored fixation dot: red and blue indicated the *Same* and *Different* conditions, respectively (Fig. 1c). On each trial, the stimulus was presented for 0.6 s, followed by a 0.2-s speed change (increase or decrease) of the attended stimulus on the target side and a 0.2-s fixation interval. Then 1 of the 7 test stimuli (0°, 5°, 10°, 15°, 345°, 350°, and 355° clockwise from the vertical) was randomly presented on the ignored side for 0.2 s. Following both stimuli subjects performed two separate responses. Subjects were first asked to make a two-alternative forced-choice (2AFC) judgment on the motion direction of the test stimulus on the ignored side, either leftward or rightward. Then, subjects made another 2AFC judgment on the speed change of the attended stimulus on the target side, either increased or decreased (Fig. 1d). Speed discrimination thresholds were determined by QUEST (75% correct)^59^ before each MAE task to ensure that subjects performed equally well in the *Same* and *Different* conditions. The change detection thresholds, response accuracy, and reaction times in the *Same* and *Different* conditions are shown in Supplementary Fig. 1a. A repeated-measures ANOVA revealed that there were no significant differences (all *p* > 0.05) in all these measurements between the two conditions.

Fig. 2a shows the psychometric functions for each of the conditions. We plotted the percentage of trials in which subjects indicated directions for the test stimuli that were opposite to the motion-direction of adapting dots on the ignored side as a function of the real motion-direction of the test stimulus. For each condition, to quantitatively measure the MAE magnitude, we fit the psychometric values at the seven test motion-directions with a cumulative normal function and interpolated the data to find the motion-direction expected to be perceived as the vertical in 50% of the trials. The MAEs of each condition are shown in Fig. 2b and were submitted to a repeated-measures ANOVA with test (pre and post) and category (*Same* and *Different*) as within-subject factors. The main effect of test (F_1,26_ = 0.213, *p* = 0.648, partial eta-squared, η_p_^2^ = 0.008) and category (F_1,26_ = 3.016, *p* = 0.094, η_p_^2^ = 0.104) was not significant, but the interaction between these two factors was significant (F_1,26_ = 7.252, *p* = 0.012, η_p_^2^ = 0.218). Therefore, for both the pre- and post-tests, we computed the global effect of MAE (*GE_MAE_*) to quantify how much the MAE increased during the *Same* condition relative to the *Different* condition. The global effect of MAE was calculated as follows:

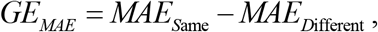

where *MAE*_*S*ame_ and *MAE*_*D*ifferent_ are the MAE of the *Same* and *Different* conditions, respectively. We hypothesized that if the global FBA effect was able to be induced by stimuli within the same category, the MAE in the *Same* condition should be significantly higher than that in the *Different* condition. The *GE_MAE_* then should be significantly higher than zero. However, if the global effect of FBA cannot be induced by the same category, the *GE_MAE_* should not be significantly different than zero. In the pre-test (before the category learning), the *Same* condition did not show a significantly higher MAE than the *Different* condition, and the *GE_MAE_* was not significantly different than zero (t_26_ = −0.052, *p* = 0.959, Cohen’s *d* = −0.020). In the post-test (after the category learning), however, the *Same* condition showed a significantly higher MAE than the *Different* condition, and the *GE_MAE_* was significantly above zero (t_26_ = 3.137, *p* = 0.004, Cohen’s *d* = 1.230) (Fig. 2c, left). These results demonstrated that a global FBA effect can be induced by the categorized features, with the MAE from adapting to the ignored stimulus significantly elevated when it fell within the same category as the attended stimulus on the target side.

**Fig. 2.**
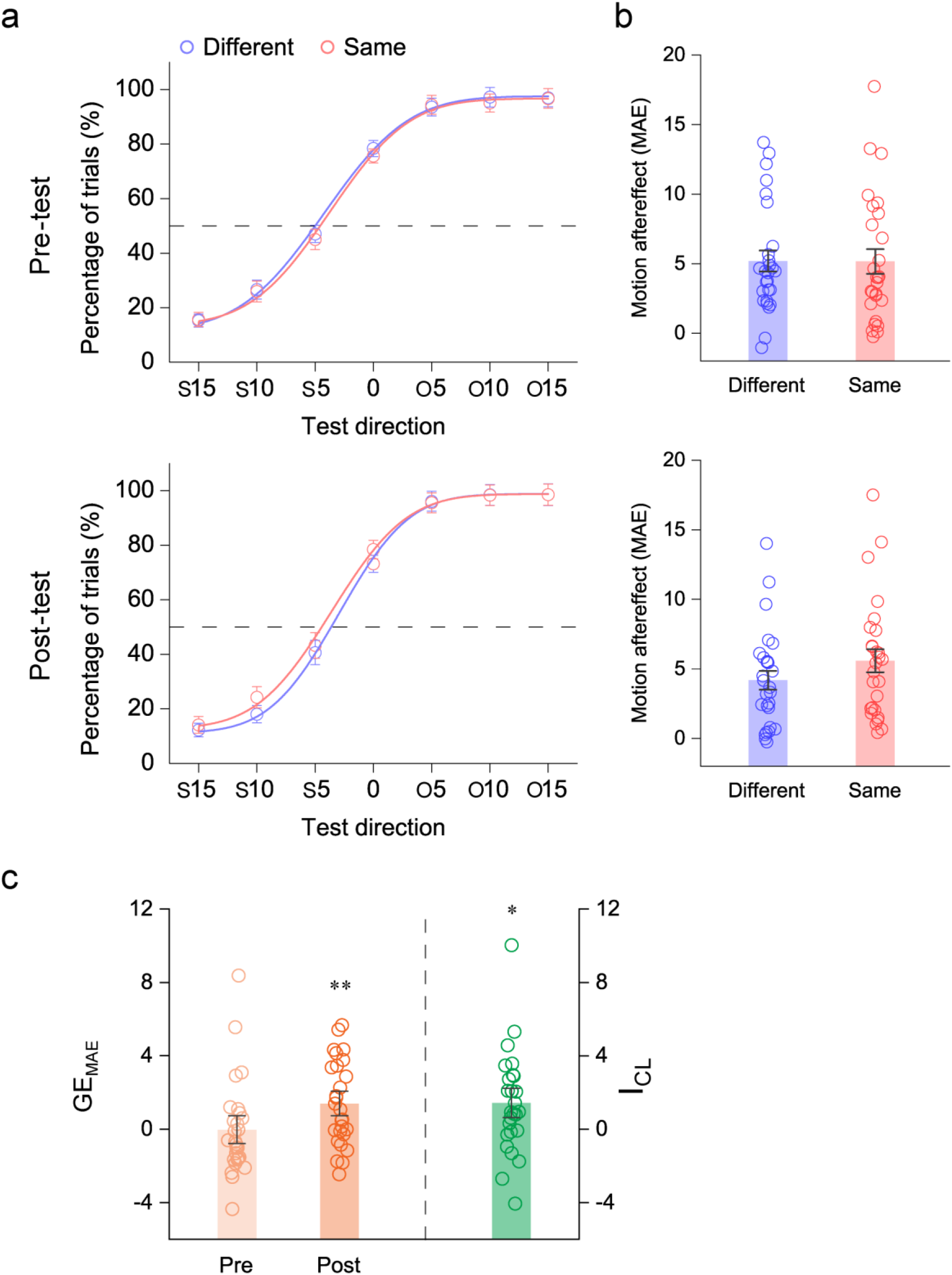
Psychophysical results. **a** Psychometric functions showing motion direction judgments of the *Same* and *Different* conditions for the pre-test (top) and post-test (bottom). Data points averaged across subjects were fit using a cumulative normal function. The abscissa refers to the seven motion directions of the test stimuli. S and O indicate that the test stimulus has the same or opposite motion direction (leftward or rightward) as the adaptor, respectively. The ordinate refers to the percentage of trials in which subjects indicated that the moving direction of a test stimulus was the opposite to the adaptor. Error bars denote 1 SEM calculated across subjects. **b** The magnitude of the motion aftereffect (MAE) in the *Same* and *Different* conditions for the pre- and post-tests. Error bars denote 1 SEM calculated across subjects and colored dots denote the data from each subject. **c** Global effect of the MAE (*GE_MAE_*) for the pre- and post-tests, and the corresponding category learning index in the MAE task (*I_CL_*). Error bars denote 1 SEM calculated across subjects and colored dots denote the data from each subject. \**p* < 0.05 and \*\**p* < 0.01, respectively.

We further calculated a category learning index (*I_CL_*) to quantify how much the global effect of MAE changed after the category learning relative to before. The learning index was calculated as follows:

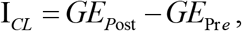

where *GE*_*p*re_ and *GE*_*P*ost_ are the global effect of MAE during the pre- and post-tests, respectively. The index quantified the global effect difference (i.e., increase) for the same category before and after category learning while subtracting out the difference for the different category. By contrasting the differences for the same and different categories, the index isolated those effects specific to the same category and the learning task and distinguished these from general practice effects or common sources of variance (e.g., day-to-day measurement variation, stimulus repetition). An index significantly above zero indicates that category learning induced the global attentional effect for stimuli from the same category. We found that the *I_CL_* was significantly greater than zero (t_26_ = 2.693, *p* = 0.012, Cohen’s *d* = 1.056, Fig. 2c, right), further demonstrating that the global effect of FBA can be induced by learned categorical features.

### fMRI FBA measurements

Before and after the category learning task, we measured BOLD signals evoked by the target and ignored sides of the display. The fMRI experiment used a block design and consisted of 6 functional runs. Each run consisted of eight stimulus blocks of 16 s, interleaved with eight blank intervals of 16 s. There were two equally common different types of stimulus blocks: *Same* and *Different*. In the *Same* condition, the category of the attended stimulus on the target side matched the category of the unattended stimulus on the ignored side (half the blocks); the *Different* condition was defined as a mismatch between categories (half the blocks) (Fig. 1e). Each stimulus block was randomly repeated four times in each run, and the attended stimulus for each stimulus block was indicated by a colored fixation dot: red and blue indicated the *Same* and *Different* conditions, respectively. Each stimulus block consisted of 8 trials; on each trial, the stimulus was presented for 0.8 s, followed by a fixed 1.2-s fixation interval. Subjects did a 0.2-s speeded discrimination task for the attended stimulus at threshold, measured by the QUEST staircase procedure (75% correct)^59^ before scanning to ensure that subjects would perform equally well in the *Same* and *Different* conditions. The change detection thresholds, response accuracy, and reaction times for the *Same* and *Different* conditions are shown in Supplementary Fig. 1b. A repeated-measures ANOVA revealed that there was no significant difference (all *p* > 0.05) in all these measurements between the two conditions.

### Region of interest analysis

Regions of interest (ROIs) in V1–V4 and MT+ were defined using the procedure outlined in the Methods section based on retinotopic mapping and functional localizers. BOLD signals were extracted from these ROIs and then averaged according to the category match (the *Same* and *Different* conditions) following the procedure described in the Methods section. For each stimulus block, the 2 s preceding the block served as a baseline, and the mean BOLD signal from 5 s to 16 s after stimulus onset was used as the measure of response amplitude. Similar to the psychophysical MAE test, during both the pre- and post-fMRI experiments, for each subject and each ROI, we computed the global effect of BOLD signal (*GE_BOLD_*) to quantify the magnitude of the BOLD signal increase during the *Same* condition relative to the *Different* condition in the ROI. The index was calculated as follows:

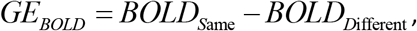

where *BOLD*_*S*ame_ and *BOLD*_*D*ifferent_ are the mean BOLD signals in the *Same* and *Different* conditions, respectively. We hypothesized that if a cortical area was sensitive to the category-induced global effect, the area would show a higher response in the *Same* condition than that in the *Different* condition, and the *GE_BOLD_* of this area would be significantly higher than zero. However, if the cortical area was not sensitive to the category-induced global effect, the *GE_BOLD_* would not be significantly different than zero. Fig. 3 shows the mean BOLD signals in the *Same* and *Different* conditions for the pre- and post-tests, and the corresponding *GE_BOLD_* of the target and ignored sides. For the target side (Fig. 3a), V1–V4 and MT+ did not show a significantly higher response in the *Same* condition than that in the *Different* condition, and none of these areas showed a *GE_BOLD_* significantly different than zero in either pre-test (V1: t_18_ = −0.704, *p* = 0.490, Cohen’s *d* = −0.332; V2: t_18_ = 0.566, *p* = 0.579, Cohen’s *d* = 0.267; V3: t_18_ = −0.142, *p* = 0.889, Cohen’s *d* = −0.067; V4: t_18_ = 0.627, *p* = 0.539, Cohen’s *d* = 0.296; MT+: t_18_ = 0.804, *p* = 0.432, Cohen’s *d* = 0.379) or post-test (V1: t_18_ = −0.179, *p* = 0.860, Cohen’s *d* = −0.084; V2: t_18_ = −0.034, *p* = 0.973, Cohen’s *d* = −0.016; V3: t_18_ = 0.778, *p* = 0.447, Cohen’s *d* = 0.367; V4: t_18_ = 1.335, *p* = 0.198, Cohen’s *d* = 0.629; MT+: t_18_ = 0.828, *p* = 0.418, Cohen’s *d* = 0.390). These findings confirmed that there was no significant difference in task difficulty or, presumably, attention between the *Same* and *Different* conditions in both pre- and post-tests. For the ignored side (Fig. 3b), none of these areas showed a significantly higher response in the *Same* condition than that in the *Different* condition, and their *GE_BOLD_* s were not significantly different than zero in the pre-test (V1: t_18_ = −0.352, *p* = 0.729, Cohen’s *d* = −0.166; V2: t_18_ = 0.394, *p* = 0.698, Cohen’s *d* = 0.186; V3: t_18_ = 0.416, *p* = 0.683, Cohen’s *d* = 0.196; V4: t_18_ = 0.879, *p* = 0.391, Cohen’s *d* = 0.414; MT+: t_18_ = −0.114, *p* = 0.910, Cohen’s *d* = −0.054). In the post-test, V1–V4 also did not show a significantly higher response in the *Same* condition than that in the *Different* condition, and none of these areas showed a *GE_BOLD_* significantly different than zero (V1: t_18_ = −1.379, *p* = 0.185, Cohen’s *d* = −0.650; V2: t_18_ = −0.708, *p* = 0.488, Cohen’s *d* = −0.334; V3: t_18_ = 1.483, *p* = 0.155, Cohen’s *d* = 0.699; V4: t_18_ = 1.197, *p* = 0.247, Cohen’s *d* = 0.564). However, MT+ showed a significantly greater response in the *Same* condition than that in the *Different* condition, and *GE_BOLD_* in this area was significantly above zero (t_18_ = 4.993, *p* < 0.001, Cohen’s *d* = 2.354), demonstrating that only MT+ showed a category-induced global effect, with responses to the ignored stimulus significantly elevated when it shared the same category as the attended stimulus.

**Fig. 3.**
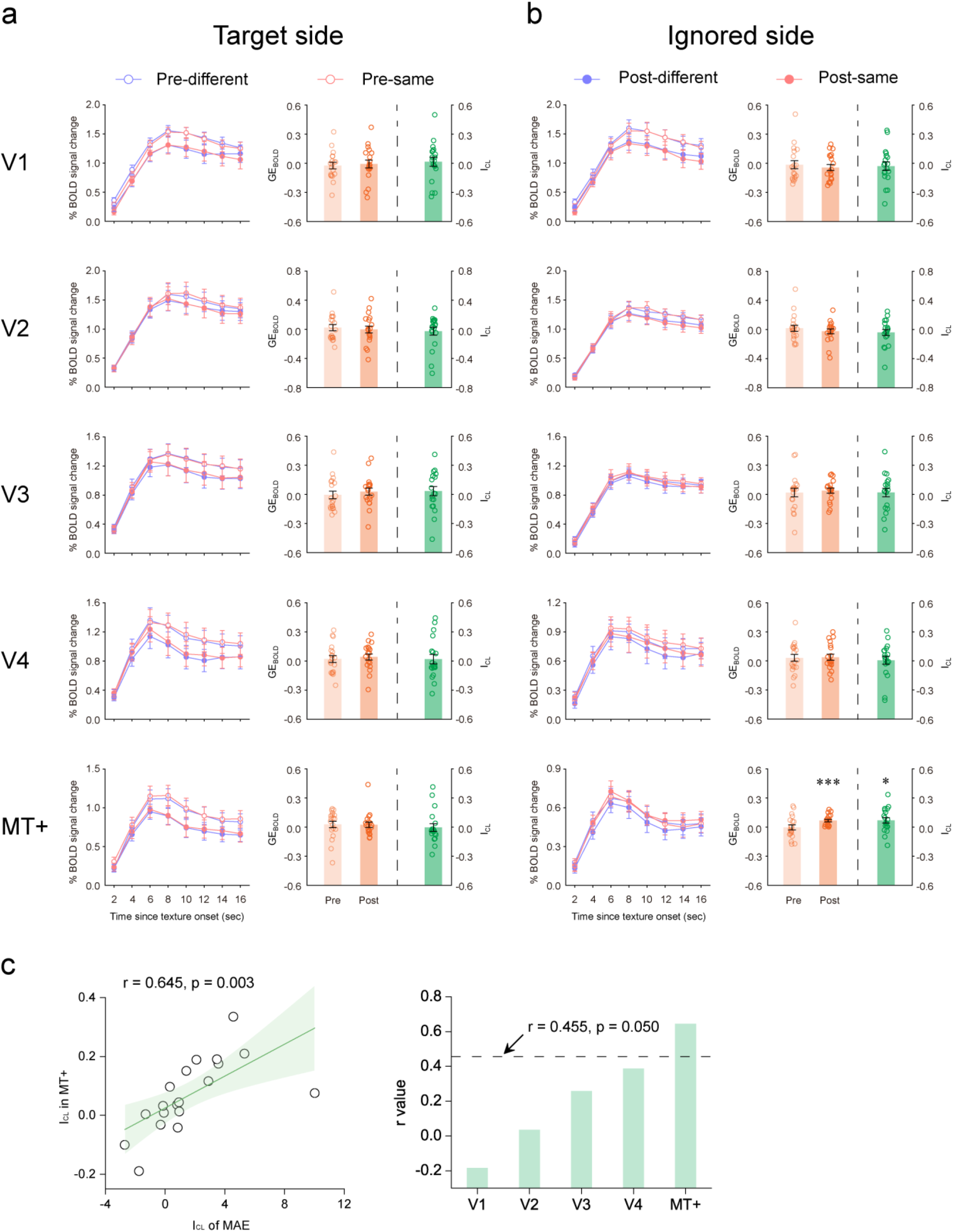
Results of ROI analyses. **a** fMRI ROI analyses for the target side. The graphs on the left side of show the BOLD signals averaged across subjects in the V1–V4 and MT+ ROIs during the *Same* (red lines) and *Different* (blue lines) conditions for the pre- (open circles) and post- (solid circles) tests. Error bars denote 1 SEM calculated across subjects at each time point. The graphs on the right side show the global effect of the BOLD amplitude (*GE_BOLD_*: difference in the BOLD signal between the *Same* and *Different* conditions) in V1–V4 and MT+ in the pre- and post-tests, and the corresponding difference measure *I_CL_* for the BOLD response. Error bars denote 1 SEM calculated across subjects and colored dots denote the data from each subject. **b** fMRI ROI analyses for the ignored side. See caption for (**a**) for a description of each type of graph (\**p* < 0.05 and \*\*\**p* < 0.001, respectively). **c** Left: Correlations between the behavioral *I_CL_* of MAE in psychophysical experiments and the neural *I_CL_* of BOLD amplitude in MT+ across individual subjects. Right: Correlation coefficients (r values) between *I_CL_* of MAE and the *I_CL_* of BOLD amplitude in other cortical visual areas across individual subjects.

Similar to the psychophysical MAE test, for each subject and each ROI, we calculated a category learning index (*I_CL_*) based on the BOLD measures to quantify how much the global effect changed after category learning relative to before. An index significantly above zero would indicate that category learning induced a global effect to the same category. We found that the *I_CL_* was significantly greater than zero in MT+ of the ignored side (t_18_ = 2.450, *p* = 0.025, Cohen’s *d* = 1.155), but not in V1–V4 of the ignored side (all t_18_ < 0.982, *p* > 0.339, Cohen’s *d* < 0.463, Fig. 3b), and not in V1-MT+ of the target side (all t_18_ < 0.709, *p* > 0.487, Cohen’s *d* < 0.334, Fig. 3a), further confirming that the global effect of FBA can be induced by the categorized features in MT+ only. In addition, to evaluate further the role of MT+ activity in the category-induced global effect of FBA, we calculated correlation coefficients between our psychophysical and fMRI measures on the ignored side across individual subjects. The *I_CL_* of MAE was significantly correlated with the *I_CL_* of BOLD amplitude in MT+ (r = 0.645, *p* = 0.003, Fig. 3c, left), but not in V1–V4 (all r < 0.388, *p* > 0.101, Fig. 3c, right). These results further indicate a close relationship between the category-induced global MAE effect and MT+ activity.

### Whole-brain and correlation analyses

To identify additional cortical or subcortical area(s) that might show a category-induced global effect similar to that found in MT+, we performed a whole-brain group analysis. We used a whole-brain search with a general linear model (GLM) procedure^60^ to identify brain regions where the *GE*_*P*ost_ was significantly greater than the *GE*_Pre_ (i.e., the *I_CL_* for the BOLD response was significantly above zero). Statistical maps were thresholded at *p* < 0.05 and corrected for multiple comparisons using the false discovery rate (FDR) correction^61^. Results showed that regions of the posterior parietal cortex (PPC, t_18_ = 2.922, *p* = 0.009, Cohen’s *d* = 1.377), anterior cingulate cortex (ACC, t_18_ = 3.486, *p* = 0.003, Cohen’s *d* = 1.643), inferior frontal junction (IFJ, t_18_ = 3.391, *p* = 0.003, Cohen’s *d* = 1.599), and the dorsolateral prefrontal cortex (DLPFC, t_18_ = 3.556, *p* = 0.002, Cohen’s *d* = 1.676) demonstrated a greater global effect in the post-test than that in the pre-test (Fig. 4a). We also analyzed each area’s *GE_BOLD_* separately in both the pre-and post-tests. All of these areas showed a significantly higher response in the *Same* condition than that in the *Different* condition and their *GE_BOLD_* s were significantly above zero in the post-test (all t_18_ > 3.199, *p* < 0.005, Cohen’s *d* > 1.508), but not in the pre-test (all t_18_ < 1.544, *p* > 0.140, Cohen’s *d* < 0.728) (Fig. 4c). Furthermore, we calculated the correlation coefficients between the *I_CL_* in MT+ and that in these cortical areas across individual subjects. Results showed that the *I_Œ_* in MT+ correlated significantly with that in IFJ (r = 0.581, *p* = 0.009), but not with that in PPC (r = −0.088, *p* = 0.721), ACC (r = −0.281, *p* = 0.243), or DLPFC (r = −0.099, *p* = 0.686) (Fig. 4d). More importantly, this correlation coefficient in IFJ was significantly larger than those in PPC (t_18_ = 2.163, *p* = 0.015, Cohen’s *d* = 0.992), ACC (t_18_ = 2.585, *p* = 0.005, Cohen’s *d* = 1.186), and DLPFC (t_18_ = 2.390, *p* = 0.008, Cohen’s *d* = 1.097). Altogether, our results suggest that the category-induced global FBA effect in MT+ might derive from feedback projections from IFJ.

**Fig. 4.**
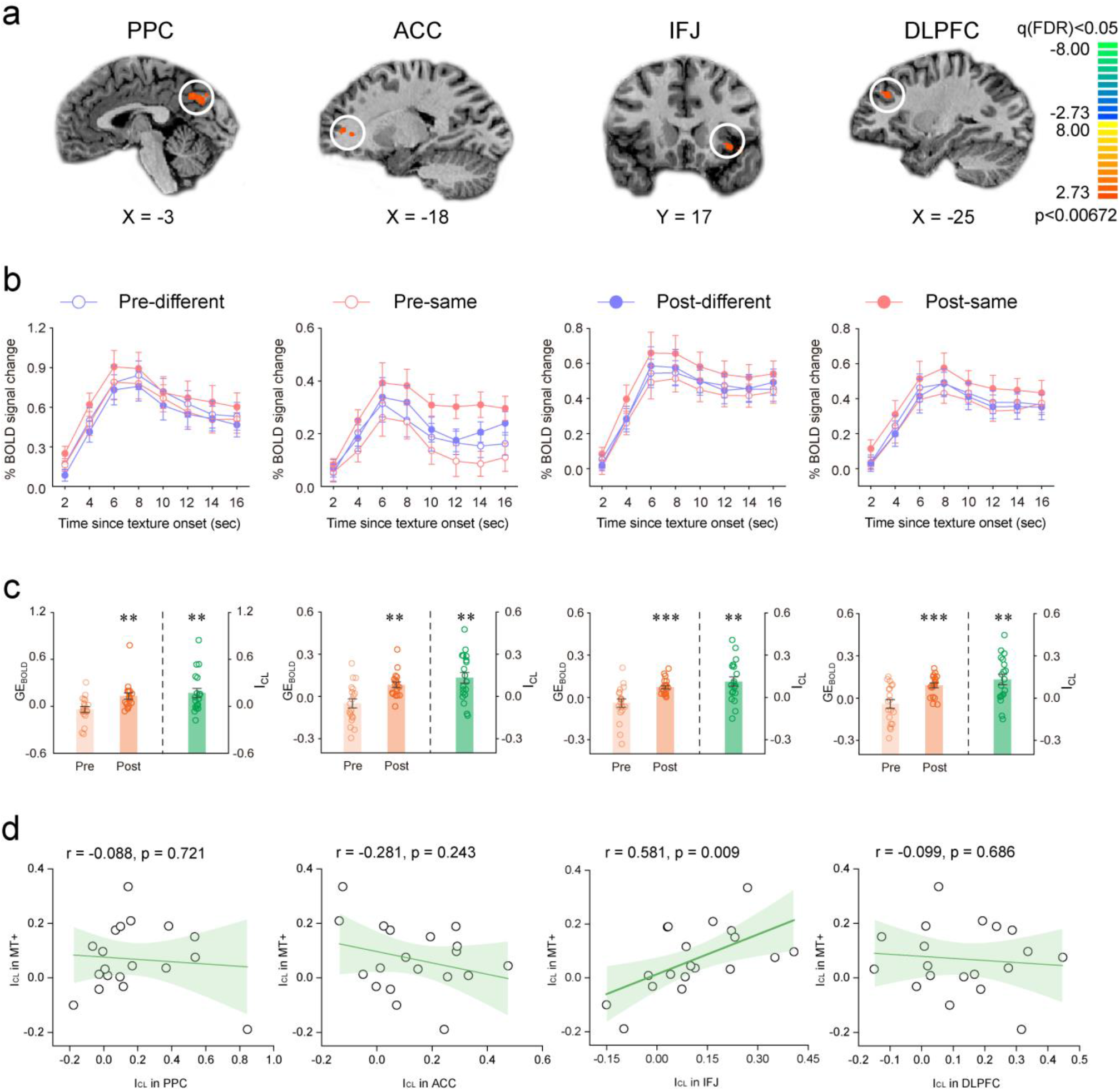
Results of whole-brain and correlation analyses. **a** Areas identified in the whole-brain analysis: PPC, ACC, IFJ, and DLPFC. All showed a significantly greater difference between the *Same* and *Different* conditions in the post-test (*GE*_*P*ost_) than that in the pre-test (*GE*_Pre_) (i.e., the neural *I_CL_* measure was significantly above zero). Statistical maps were thresholded at *p* < 0.05 and corrected for multiple comparisons using the false discovery rate correction (q < 0.05). **b** BOLD signals averaged across subjects within each corresponding ROI during the *Same* (red) and *Different* (blue) conditions for the pre- (open circles) and post- (solid circles) tests. Error bars denote 1 SEM calculated across subjects at each time point. **c** *GE_BOLD_* for each ROI in the pre- and post-tests, and the corresponding neural *I_CL_*. Error bars denote 1 SEM calculated across subjects and colored dots denote the data from each subject. \*\**p* < 0.01 and \*\*\**p* < 0.001, respectively. **d** Correlations between the neural in MT+ and that in each ROI across individual subjects. Shaded area represents 95% confidence intervals for each correlation.

### Effective connectivity analysis

To directly confirm whether the category-induced global FBA effect in MT+ was supported by feedback from IFJ, we performed dynamic causal modeling (DCM) analysis^62^ in SPM12 to examine functional changes in interregional connectivity related to the *Same* condition. Given the extrinsic visual input into both TMT+ (the ROI in MT+ evoked by the stimulus on the target side) and IMT+ (the ROI in MT+ evoked by the stimulus on the ignored side), we posited a network of bidirectional connections linking PPC, ACC, IFJ, DLPFC, TMT+, and IMT+, and assessed the change in the modulatory effect by the *Same* condition between the pre- and post-tests (Fig. 5a). Results showed that, relative to the pre-test, the modulatory input (the *Same* condition) significantly increased the feedback from IFJ to IMT+ (t_18_ = 2.301, *p* = 0.034, Cohen’s *d* = 1.056) and (marginally) significantly increased the connection from IFJ to DLPFC (t_18_ = 1.859, *p* = 0.079, Cohen’s *d* = 0.853) and decreased the connection from PPC to ACC (t_18_ = −1.912, *p* = 0.072, Cohen’s *d* = −0.877). These results support the idea that the category-induced global FBA effect in MT+ is due to feedback from IFJ rather than from PPC, ACC, or DLPFC (Fig. 5b). To further evaluate the role of these modulatory effects in the category-induced global FBA effect, across individual subjects, we calculated the correlation coefficients between the modulatory effect difference and *I_CL_* in MT+, and between the modulatory effect difference and *I_CL_* of MAE from the psychophysical experiments. Results showed that both the *I_CL_* in MT+ (r = 0.474, *p* = 0.040, Fig. 5c) and *I_CL_* of MAE (r = 0.498, *p* = 0.030, Fig. 5d) correlated significantly with the modulatory effect difference between the pre- and post-tests in IFJ, but not in PPC (*I_CL_* in MT+: r = 0.047, *p* = 0.850; *I_CL_* of MAE: r = 0.244, *p* = 0.315), ACC (*I_CL_* in MT+: r = 0.080, *p* = 0.746; *I_CL_* of MAE: r = −0.058, *p* = 0.815), or DLPFC (*I_CL_* in MT+: r = −0.085, *p* = 0.729; *I_CL_* of MAE: r = −0.150, *p* = 0.540), further suggesting that IFJ may be the source of the category-induced global FBA effect in the human visual system.

**Fig. 5.**
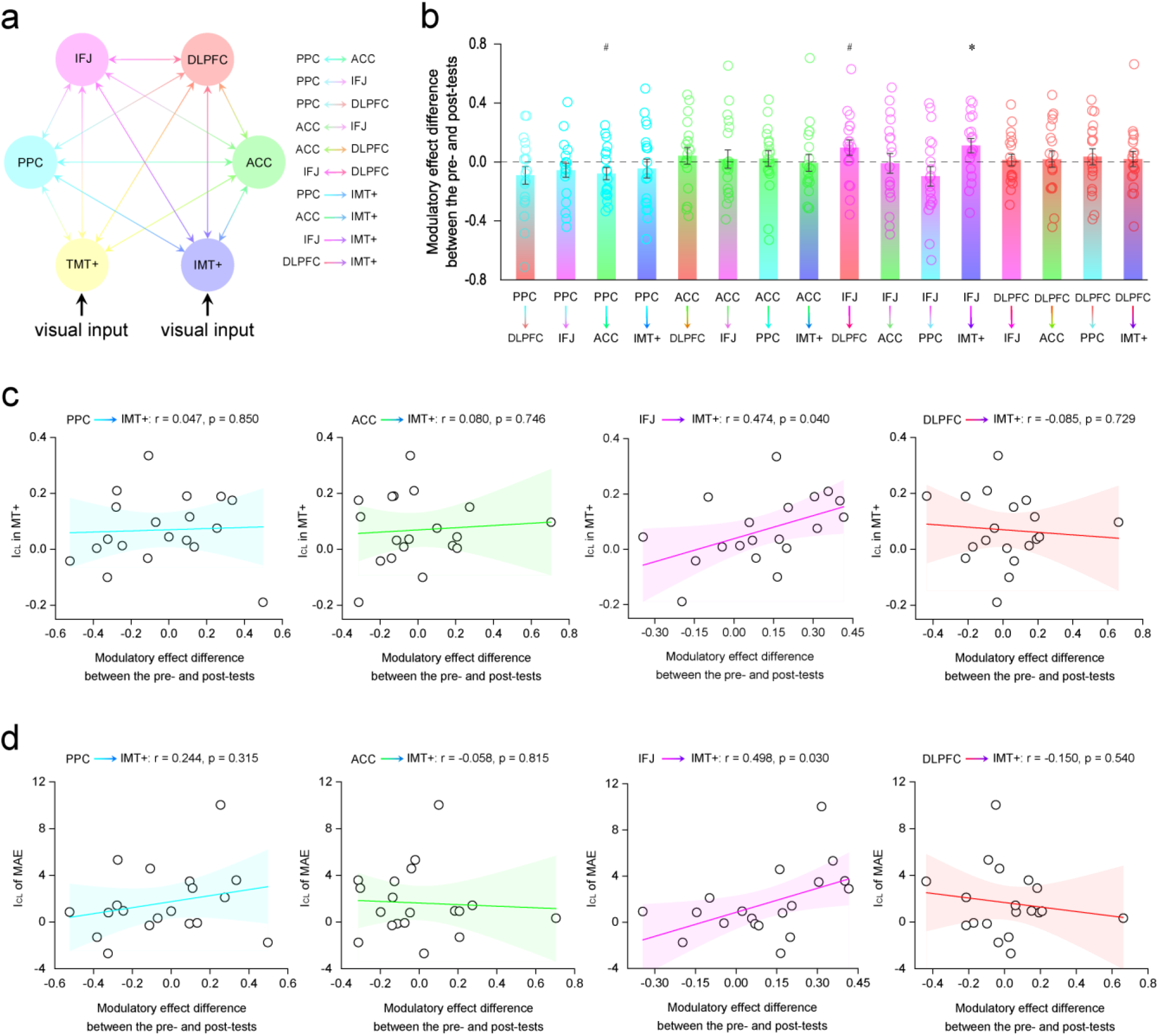
Results of DCM analysis. **a** Given the extrinsic visual input into both TMT+ (the ROI in MT+ evoked by the stimulus on the target side) and IMT+ (the ROI in MT+ evoked by the stimulus on the ignored side), bidirectional connections were hypothesized to exist among the PPC, ACC, IFJ, DLPFC, TMT+, and IMT+. The modulatory input (the *Same* condition) was hypothesized to affect the bidirectional connections between PPC and ACC, between PPC and IFJ, between PPC and DLPFC, between ACC and IFJ, between ACC and DLPFC, and between IFJ and DLPFC, as well as the intrinsic connections from PPC, from ACC, from IFJ, and from DLPFC to IMT+. **b** Differences in modulatory effects between the pre- and post-tests and their significance levels (#0.05 < *p* < 0.079; \**p* < 0.05). Error bars denote 1 SEM calculated across subjects and colored dots denote the data from each subject. **c** Correlations between the neural *I_CL_* in MT+ and the modulatory effect difference between the pre- and post-tests in PPC, ACC, IFJ, and DLPFC across individual subjects. Shaded area represents 95% confidence intervals for each correlation. **d** Correlations between the behavioral *I_CL_* of MAE in psychophysical experiments and the modulatory effect difference between the pre- and post-tests in PPC, ACC, IFJ, and DLPFC across individual subjects. Shaded area represents 95% confidence intervals for each correlation.

## Discussion

Although the global FBA effect, based on the “feature-similarity gain model”^21^, has been studied extensively and intensively for many years, visual perception in our daily life is often categorical and the ecological importance of a stimulus is usually determined by the category to which the stimulus belongs. Thus, researchers have questioned whether the global FBA effect can be generalized beyond the same stimuli in an experimental setting and applied to our daily life. We trained human subjects to classify motion directions into two discrete categories and found that after training, the attentional effect of a motion direction globally spread to the unattended motion directions within the same category as the attended motion direction but not to motion directions from the different category, in both behavior (Fig. 2) and fMRI (Fig. 3). Note that the attended motion direction in the same category and that in the different category were both 25° away from the unattended motion direction (i.e., they had the equal feature similarity, Fig. 1c), therefore the global FBA effect in our study was induced by the feature categorization rather than similarity. Intriguingly, several studies have also demonstrated that FBA does not rely on feature similarity in primate prefrontal cortex^63^ and does not reflect a monotonic similarity between attended and preferred feature, but rather shows a non-monotonic function^16,64,65^ that coincides with the feature category boundary^66^. Furthermore, two recent studies have demonstrated that attention to high-level object categories (human faces and bodies) show similar global effects of attention to simple individual low-level features^67,68^. Our results extend these findings: the category-induced global FBA effect evident in our study further suggests that FBA indeed plays an important role in adapting to or surviving the natural environment. In a cluttered scene where features are categorically organized, relative to the feature similarity, feature categorization may be advantageous to globally spread FBA from one stimulus within a category to other stimuli in the same category.

Our study not only revealed a category-induced global FBA effect but also identified the IFJ as a source of this effect in the human visual system. First, IFJ responses showed a category-induced global FBA effect, with neural responses to the ignored stimulus significantly elevated when it shared the same category as the attended stimulus (Fig. 4c) and this response correlated significantly with the category-induced global effect in MT+ (Fig. 4d). Second, the DCM analysis indicated that the category-induced global effect in MT+ was derived by feedback from IFJ rather than PPC, ACC, or DLPFC (Fig. 5b). Moreover, this increased feedback from IFJ significantly predicted the category-induced global effect in the MT+ (Fig. 5c) and the MAE in the psychophysical experiment (Fig. 5d). In addition, these results cannot be explained by eye movements, attention shifting, or task difficulty, as none of factors resulted in any significant differences between our conditions (Supplementary Fig. 2).

The IFJ is a region ventrolateral to FEF and is anatomically localized at the intersection of the precentral sulcus and the inferior frontal sulcus^69,70^. Several studies have shown that IFJ has anatomical connections with sensory, parietal, and prefrontal areas^71,72^ and functionally interacts with both ventral and dorsal cortical brain structures^73–75^. The IFJ is therefore ideally suited to act as a neural system important for many different cognitive processes, such as the FBA and categorization. Indeed, using neurophysiological and brain imaging techniques, a number of previous studies have demonstrated that the IFJ not only plays a key role in the feature similarity-based global FBA effect^18,26,28,33,48,76–78^ but also strongly reflects the representation of category membership^79–86^. Our current results support and extend the function of IFJ in both of these two processes. A crucial involvement of IFJ in both the feature similarity- and categorization-based global FBA effects indicates that IFJ can direct top-down control to both sensory and cognitive processes within posterior cortices, consistent with previous studies showing flexible cognitive function in prefrontal cortex^87^. While it remains to be examined whether the feature similarity- and categorization-based global FBA effects are mediated by the same or different subpopulations of IFJ neurons, a central role for this area in the FBA resonates well with the proposal that the IFJ is a critical neural substrate that underlies the control of attention and awareness in general. Moreover, our results indicate that the IFJ is able to represent categories not only at an attended location but also in the opposite visual hemifield, a finding that is consistent with recent data suggesting that category learning effect can transfer to the opposite visual hemifield ^88^ (but see^40^). Taking into account the generalized function of the IFJ across both the FBA and categorization, and our fMRI findings, we speculate that attending to a feature from a category may elevate the neural response to this category in the IFJ and thus actively and globally result in top-down biasing signals to other features falling within the same category (Fig. 6). Identifying IFJ as a source of the category-induced global effect for FBA and the corresponding speculation here derive mainly from our DCM analyses, which depends on timeseries models of fMRI data for an interpretation of causality^62^. Although this interpretation of causality in our study finds support in previous lesion^26,28^ and transcranial magnetic stimulation^77,89^ studies showing a causal effect of prefrontal cortical areas in controlling FBA, given the limitation of fMRI in our study, further work is needed using neurophysiological techniques to parse the crucial involvement of prefrontal cortical areas in the category-induced global effect of FBA.

**Fig. 6.**
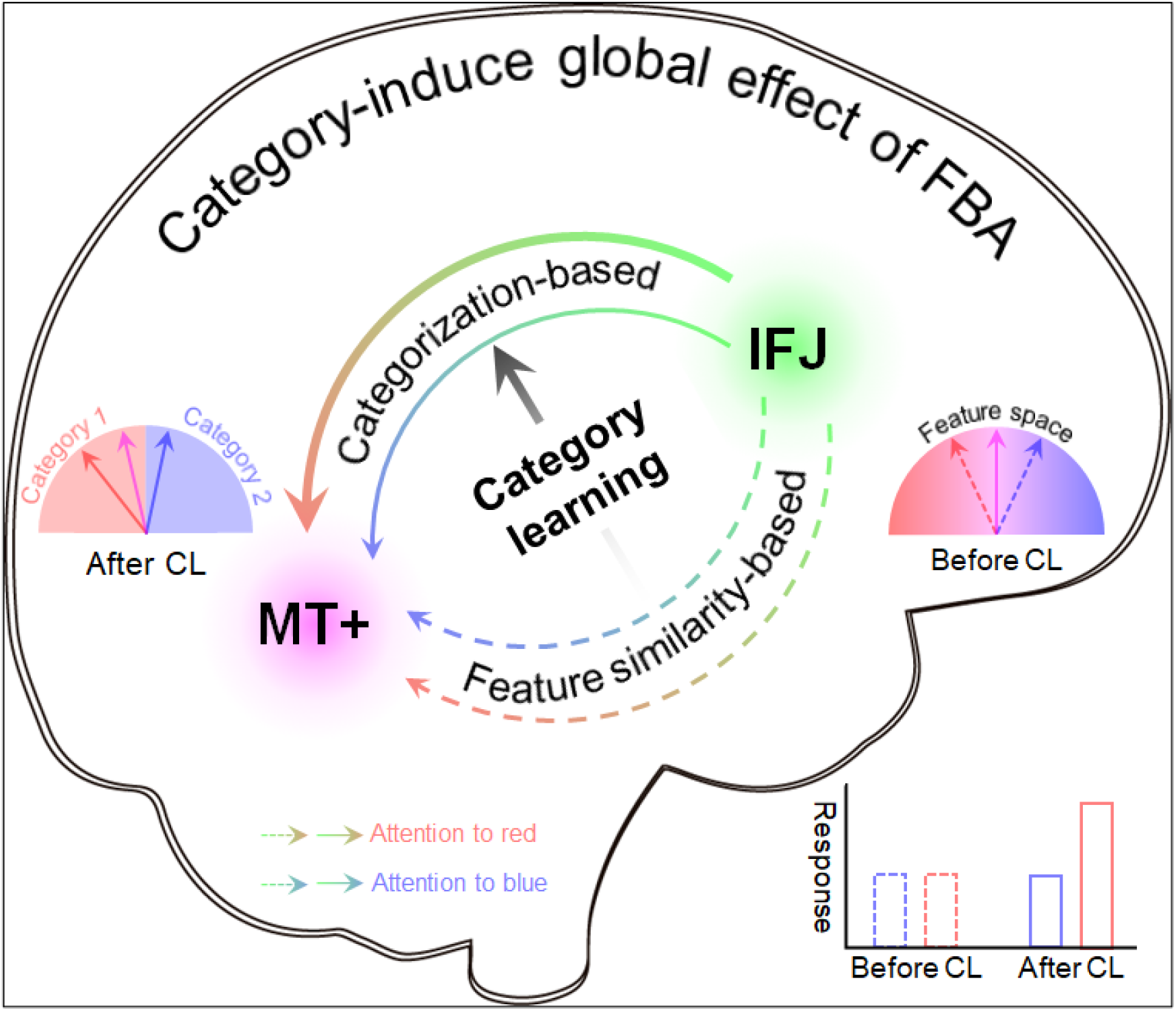
Schematic illustration of the category-induced global FBA effect. Before category learning (see right side of the image and dashed lines) the angular difference in feature space between the unattended stimulus (purple arrow) and each of the two possible attended stimuli (red and blue arrows) are equal. Therefore, the top-down feedback from IFJ to MT+ (dashed arrows) and neural responses to the unattended stimulus in MT+ (dashed bars) are increased equally when attending to either of the attended stimuli, resulting in no significant differences in the feature similarity-based global FBA effect between two conditions. In contrast, after category learning (see left side of the image and solid lines) the unattended stimulus and one of the attended stimuli (red arrow) are categorized as belonging to the same category (Category 1), whereas the other attended stimulus (blue arrow) belongs to another category (Category 2). Attending to the stimulus from category 1 (red arrow) elevates the neural response to the entire category in IFJ and actively increases top-down biasing signals to MT+ for all category members including the unattended stimulus (thick-solid arrows). This results in a greater top-down biasing signal for same category than different category stimuli, resulting in a significant category-induced global FBA effect (solid bars in graph on the bottom right).

Intriguingly, our study did not find category-induced global FBA effects in V1–V4, in disagreement with previous EEG^17,29,30^, MEG^11,31^, and fMRI^20,32,33^ studies reporting the feature similarity-based global FBA effects in early visual cortex. One possible explanation is that neurons throughout the early visual cortex are known to select different feature dimensions^90,91^, but less so for the representation of category membership. Indeed, virtually all theories of category learning^35,92^ and previous neurophysiological studies^36,43,47,51,81,93^ have indicated that the representation of category membership is removed from visual characteristics of the display and primarily occurs in higher-level visual areas (rather than low-level visual areas, but see^40^), which is therefore immune to the equal feature similarity between the *Same* and *Different* conditions here (Fig. 1c). Alternatively, the top-down feedback from IFJ primarily modulates the respective motion processing areas (MT+) but not early visual cortex^94^ or decreases gradually from MT+ to early visual cortex^95,96^. Given these inconsistent findings, further work is thus needed using both feature similarity- and categorization-based paradigms to parse the crucial involvement of early visual cortex in the global modulation of FBA.

In sum, our study provides, to the best of our knowledge, the first experimental evidence supporting a category-induced global FBA effect and identifies the IFJ as the source of this effect in human visual system (Fig. 6). Extending the function of human prefrontal cortex to the category-induced global FBA effect in our study gives insight into how prefrontal cortical areas directly top-down control the attentional selection and awareness within posterior cortices^97,98^ and in real-life situations, how they flexibly deal with a multitude of cognitive processes, ranging from low-level feature similarity to high-level categorization. Combining our results with earlier studies showing the classical feature similarity-based global effects, we conclude that in our daily life, FBA may occur in a more generalized way and may be of more ecological and adaptive significances than has been previously thought.

## Methods

### Subjects

A total of 27 human subjects (10 male, 18-25 years old) were involved in the study. All of them participated in the psychophysical experiment. Twenty of them participated in the fMRI experiment but one subject in the fMRI experiment was excluded because of large head motion (>3 mm). They were naïve to the purpose of the study. They were right-handed, reported normal or corrected-to-normal vision, and had no known neurological or visual disorders. They gave written, informed consent, and our procedures and protocols were approved by the human subjects review committee of School of Psychology at South China Normal University.

### Stimuli

Visual stimuli were random dot kinetograms (RDKs), presented against a gray background (luminance: 11.55 cd/m^2^). All dots (dot diameter: 0.13°; dot luminance: 76.8 cd/m^2^; dot density: 2.0/(°)^2^; speed: 10.0°/sec) in each RDK moved in the same direction (100% motion coherence). The category boundary motion-direction was randomly selected from 17.5°, 42.5°, 317.5°, and 342.5° clockwise from the vertical for each subject. Two groups of 3 motion-directions were separated by each category boundary that centered the range of the motion-directions. As shown in Fig. 1b (the category boundary motion-direction here was 42.5°, as an example), the 6 motion-directions were evenly spaced (25° apart). The stimulus display in both psychophysical and fMRI experiments was composed of two circular regions (diameter: 8.0°) in the upper visual field (centered 8.5° to the left and right of the central fixation point). For each subject, one of these regions was attended (the target side) and the other was unattended (the ignored side) (Fig. 1c). The two sides were counterbalanced across subjects. On the category learning task (Fig. 1b) of psychophysical experiments, the dots were always presented on the target side and moved in 1 of 6 evenly spaced directions. On both the threshold measurement and motion aftereffect (MAE) tasks of psychophysical experiments (Fig. 1d) and in the fMRI experiments (Fig. 1e), the ignored side was a single field of dots moving as an adapting motion-direction (−12.5° away from the category boundary, i.e., the adaptor). The target side was comprised of two overlapping non-adapting moving dot directions: one belonged to the adapting category (−25° from the adapting motion-direction, i.e., the same category; subjects attended to these moving dots indicated the *Same* condition) and the other one belonged to the non-adapting category (25° from the adapting motion-direction, i.e., the different category; subjects attended to these moving dots indicated the *Different* condition) (Fig. 1c). In the MAE task, the test stimuli were always presented on the ignored side with one of seven motion-directions: 0°, 5°, 10°, 15°, 345°, 350°, and 355° clockwise from the vertical (Fig. 1d). For all stimuli in the current study, to minimize the possibility that subjects could focus on a single dot, half of the dots disappeared and were replaced by new dots at different random locations every 100 ms.

### Psychophysical experiments overview

The psychophysical experiments included three different tasks: the category learning, threshold measurement, and MAE. For all experiments visual stimuli were displayed on an IIYAMA color graphic monitor (model: HM204DT; refresh rate: 60 Hz; resolution: 1,280 × 1,024; size: 22 inches) at a viewing distance of 57 cm. Subjects’ head position was stabilized using a chin rest. A yellow fixation cross was always present at the center of the monitor.

### Category learning task

In the category learning task, we trained human subjects to classify motion directions into two discrete categories on the target side. As shown in Fig. 1b, each trial started with a fixation interval. The first sample stimulus (the dots moved in 1 of 6 evenly spaced directions) was presented for 800 ms, followed by a 200 ms fixation interval. Then, the second sample stimulus (the dots moved in 1 of the remaining 5 motion-directions) was presented for 800 ms and was followed by a response interval. Subjects were asked to press one of two buttons to indicate whether the second sample stimulus was in the same or different category as the first sample stimulus and received auditory feedback if their response was incorrect. The category learning task, during each day, consisted of five blocks. Each block had 120 trials, randomly interleaving 20 trials from each of the 6 motion-directions. Training was terminated when subjects’ accuracy was higher than 85% for each of the motion-directions. It took around 3~5 days (about 18.0 ± 1.31 blocks) for each subject to complete the categorization training.

### Motion aftereffect task

To examine whether the global effect of FBA can be induced by the same category, before and after the category learning task, we measured each subject’s motion aftereffects (MAEs) resulting from adapting to the moving dots on the ignored side. The MAE task consisted of two sessions (the *Same* and *Different* conditions), with the two sessions occurring on different days; the order of the two sessions was counterbalanced across subjects. For each session, subjects attended one direction of motion (the attended stimulus) within a display of two overlapping non-adapting dots on the target side, and ignored the adaptor (the unattended stimulus) on the ignored side. The attended stimulus on the target side was indicated by a colored fixation dot: red and blue indicated the *Same* and *Different* conditions, respectively (Fig. 1c). Each session consisted of 10 blocks; each block had 56 trials, from randomly interleaving 8 trials from each of the 7 test stimuli. Motion-direction of the test stimuli varied from trial to trial in randomly shuffled order, and test stimuli were presented briefly to avoid any possible dependence of attentional state on the stimulus motion-direction. Each trial began with the central fixation. The stimulus was presented for 0.6 s, followed by a 0.2-s speed change (increase or decrease) of the attended stimulus on the target side and a 0.2-s fixation interval. Then 1 of the 7 test stimuli (0°, 5°, 10°, 15°, 345°, 350°, and 355° clockwise from the vertical) was randomly presented on the ignored side for 0.2 s, followed by two response intervals. Subjects were first asked to make a two-alternative forced-choice (2AFC) judgment on the motion direction of the test stimulus on the ignored side, either leftward or rightward. Then, subjects needed to make another 2AFC judgment on the speed change (the speed discrimination threshold was measured by the threshold measurement task, see the threshold measurement task) of the attended stimulus on the target side, either increased or decreased (Fig. 1d).

### Threshold measurement task

Before each MAE task, to ensure that subjects performed equally well for the *Same* and *Different* conditions, we measured the speed discrimination thresholds of the attended stimulus on the target side with a 2AFC QUEST staircase procedure (75% correct)^59^. The stimuli and procedure of the threshold measurement task were the same as those of the MAE task, except that no test stimuli were presented. The threshold measurement task also consisted of two sessions (the *Same* and *Different* conditions), with the two sessions occurring on different days; the order of the two sessions was counterbalanced across subjects. Each session consisted of 20 QUEST staircases of 40 trials. Each trial began with the central fixation. The stimulus was presented for 0.6 s, followed by a 0.2-s speed change (increase or decrease) of the attended stimulus on the target side and a response interval. Subjects were asked to press one of two buttons to indicate the speed change of the attended stimulus and received auditory feedback if their response was incorrect. The speed of the attended stimulus varied trial by trial and was controlled by the QUEST staircase to estimate subjects’ speed discrimination threshold (75% correct).

### Psychophysical data analysis: MAE task

We first constructed a psychometric function for each condition shown in Fig. 2a. We plotted the percentage of trials in which subjects indicated directions for the test stimuli that were opposite to the motion-direction of adapting dots on the ignored side as a function of the real motion-direction of the test stimulus. For each subject and each condition, the psychometric values at the seven test motion-directions were fit with a cumulative normal function and we interpolated the data to find the motion-direction (the MAE) expected to be perceived as the vertical in 50% of the trials. On both the pre- and post-MAE tasks, we computed the global effect of MAE (*GE_MAE_*) to quantify how much the MAE increased during the *Same* condition relative to the *Different* condition. The global effect of MAE was calculated as follows:

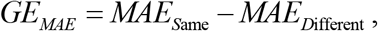

where *MAE*_*S*ame_ and *MAE*_*D*ifferent_ are the MAE of the *Same* and *Different* conditions, respectively. We hypothesized that if the global effect of FBA was able to be induced by the same category, the MAE in the *Same* condition should be significantly higher than that in the *Different* condition. The *GE_MAE_* then should be significantly higher than zero. However, if the global effect of FBA couldn’t be induced by the same category, the *GE_MAE_* should not be significantly different than zero. Moreover, we further calculated a category learning index (*I_CL_*) to quantify how much the global effect changed after the category learning relative to before. The learning index was calculated as follows:

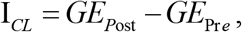

where *GE*_Pre_ and *GE*_*P*ost_ are the global effect of MAE during the pre- and post-tests, respectively. The index quantified the global effect difference (i.e., increase) for the same category before and after category learning while subtracting out the difference for the different category. By contrasting the differences for the same and different categories, the index isolated those effects specific to the same category and the learning task and distinguished these from general practice effects or common sources of variance (e.g., day-to-day measurement variation, stimulus repetition). The larger the index, the greater the category learning effect.

### fMRI measurements of FBA

Before and after the category learning task, using a block design, the fMRI experiment consisted of 6 functional runs. Each run consisted of eight stimulus blocks of 16 s, interleaved with eight blank intervals of 16 s. There were two different stimulus blocks: *Same* and *Different* conditions. In the *Same* condition, the category of attended stimulus on the target side matched the category of the stimulus on the ignored side (half the blocks); a *Different* condition was defined as a mismatch (half the blocks) (Fig.1e). Each stimulus block was randomly repeated four times in each run, and the attended stimulus in each stimulus block was indicated by a colored fixation dot: red and blue indicated the *Same* and *Different* conditions, respectively. Each stimulus block consisted of 8 trials; on each trial, the stimulus was presented for 0.8 s, followed by a fixed 1.2-s fixation interval, and subjects did a 0.2-s speed discrimination task of the attended stimulus at threshold, measured by the QUEST staircase procedure (75% correct)^59^ before scanning to ensure that subjects performed equally well for the *Same* and *Different* conditions.

### MRI data acquisition

MRI data were collected using a 3T Siemens Prisma^fit^ scanner with a 64-channel phased-array head/neck receiver coil at the Center for MRI Research at South China Normal University. In the scanner, the stimuli were back-projected via a video projector (refresh rate: 60 Hz; spatial resolution: 1,024 × 768) onto a translucent screen placed inside the scanner bore. Subjects viewed the stimuli through a mirror located above their eyes. The viewing distance was 90 cm. Blood oxygen level-dependent (BOLD) signals were measured with an echo-planar imaging sequence (TE: 30 ms; TR: 2,000 ms; FOV: 196 × 196 mm^2^; matrix: 112 × 112; flip angle: 90°; slice thickness: 2 mm; gap: 1 mm; number of slices: 58, slice orientation: axial). A high-resolution 3D structural data set (3D MPRAGE; 0.5 × 0.5 × 1 mm^3^ resolution; TR: 2,530 ms; TE: 1.94 ms; FOV: 256 × 256 mm^2^; flip angle: 7°; number of slices: 176; slice orientation: sagittal) was collected in the same session before the functional scans. Subjects underwent three sessions, one for the retinotopic mapping and ROI localization, and the other two for the main experiment.

### MRI data analysis

The MRI data analysis, ROI analysis, whole-brain group analysis, and DCM of this study closely followed those used in our previous studies^33,99–101^ and therefore, for consistency, we largely reproduce that description here, noting differences as necessary.

### FMRI preprocessing

The anatomical volume collected from each subject in the retinotopic mapping session was transformed into a brain space that was common for all subjects^102^ and then inflated using BrainVoyager QX. Functional volumes in all sessions for each subject were preprocessed, including 3D motion correction, linear trend removal, and high-pass (0.015 Hz)^103^ filtering using BrainVoyager QX. Head motion within any fMRI session was < 3 mm for all subjects. The images were then aligned to the anatomical volume in the retinotopic mapping session and transformed into Talairach space^102^. The first 8-s of BOLD signals were discarded to minimize transient magnetic saturation effects.

### Retinotopic mapping and visual ROI definitions

Retinotopic mapping was performed for each subject individually to identify retinotopic visual areas (V1–V4) based on fMRI scans performed in a separate session during pre-testing. Two fMRI scans were performed. The first was a scan implementing the standard phase-encoded method developed by Sereno et al.^104^ and Engel et al.^105^, in which subjects viewed rotating wedge and expanding ring stimuli that created traveling waves of neural activity in visual cortex. This scan was used to identify the boundaries between V1-V4 using standard procedures. An independent block-design scan was used to localize specific regions of interest (ROIs) within the V1–V4 boundaries and MT+ that were activated by the target and ignored stimuli (Fig. 1e). The scan consisted of 12 12-s stimulus blocks, interleaved with 12 12-s blank intervals, with two different types of stimuli: stationary dots and moving dots. In each stimulus block, subjects were asked to press one of two buttons to indicate a random luminance change (increase or decrease) of the stimulus. A general linear model (GLM) procedure was used for the ROI analysis. The ROIs falling within V1–V4 were defined as areas that responded more strongly to the stationary gray dots than to the blank screen (*p* < 0.05, corrected by FDR correction)^61^. The ROIs in MT+ were defined as areas that responded more strongly to the moving dots than to the stationary dots (*p* < 0.05, corrected by FDR correction)^61^.

### ROI analysis

The block-design BOLD signals were extracted from ROIs and then averaged according to the *Same* and *Different* conditions. For each stimulus block, the 2 s preceding the block served as a baseline, and the mean BOLD signal from 5 s to 16 s after stimulus onset was used as a measure of response amplitude. Similar to the psychophysical MAE test, during both the pre- and post-fMRI experiments, for each subject and each ROI, we computed the global effect of BOLD signal (*GE_BOLD_*) to quantify how much the BOLD signal increased during the *Same* condition relative to the *Different* condition in the ROI. The index was calculated as follows:

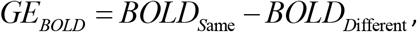

where *BOLD*_*S*ame_ and *BOLD*_*D*ifferent_ are the mean BOLD signals in the *Same* and *Different* conditions, respectively. The index is positive whenever the mean response in the *Same* condition is greater than that in the *Different* condition. The *GE_BOLD_* measures in the pre and post tests are referred to as *GE*_Pre_ and *GE*_*P*ost_, respectively. Similar to the psychophysical MAE test, for each subject and each ROI, we further calculated a neural category learning index (*I_CL_*) to quantify how much the global effect changed after category learning relative to before.

### Whole-brain group analysis

In the whole-brain group analysis, a fixed-effects general linear model (FFX-GLM)^60^ was performed for each subject on the spatially nonsmoothed functional data in Talairach space. The 1^st^-level regressors were created by convolving the onset of each stimuli block with the default BrainVoyager QX’s two-gamma hemodynamic response function. Six additional parameters resulting from 3D motion correction (x, y, z rotation and translation) were included in the model. First, for each subjects, we calculated fixed effects analyses for each *GE_BOLD_* (the *Same* condition vs. the *Different* condition) of the pre- and post-tests separately (*G*_Pre_ and *GE*_*P*ost_, respectively). Second, these FFX-GLM estimates were entered into a second-level group analysis (random-effects) of variance (t-test) and the statistical t-map was generated expressing the difference in *GE_BOLD_* between *GE*_Pre_ and *GE*_*P*ost_ (i.e., *I_CL_*). Statistical maps were thresholded at p < 0.05 and corrected by FDR correction^61^.

### Effective connectivity analysis

To directly confirm whether the category-induced global effect of FBA in MT+ was derived through the modulation of feedback from the IFJ, we applied DCM analysis^62^ in SPM12 to our fMRI data. For each subject and each hemisphere, using BrainVoyager QX, MT+ voxels were identified as those activated by the moving dots at a significance level of *p* < 0.01; all PPC, ACC, IFJ, and DLPFC voxels were identified as those activated by the stimulus block at a significance level of *p* < 0.01. The mean Talairach coordinates of MT+, PPC, ACC, IFJ, and DLPFC, and their standards errors across subjects were [−41 ± 1.26, −70 ± 1.06, −2 ± 1.79], [−25 ± 4.95, −58 ± 3.94, 38 ± 2.29], [−9 ± 1.11, 16 ± 5.02, 30 ± 2.77], [−46 ± 1.94, −1 ± 2.49, 30 ± 1.86], and [−35 ± 1.49, 33 ± 2.59, 24 ± 1.99] for the left hemisphere, and [43 ± 0.90, −64 ± 1.32, −2 ± 1.60], [25 ± 2.20, −54 ± 2.41, 43 ± 2.84], [7 ± 0.82, 21 ± 1.60, 29 ± 1.65], [43 ± 1.50, 8 ± 2.06, 25 ± 3.09], and [35 ±1.51, 36 ± 1.23, 24 ± 2.59] for the right hemisphere, respectively. For each subject and each hemisphere, these Talairach coordinates were converted to Montreal Neurological Institute (MNI) coordinates using the tal2mni conversion utility^106^. In SPM, for each of these areas, we extracted voxels within a 4-mm sphere centered on the most significant voxel and used their time series for the DCM analysis. The estimated DCM parameters were later averaged across two hemispheres using the Bayesian model averaging method^107^. DCMs have three sets of parameters: (1) extrinsic input into one or more regions; (2) intrinsic connectivities among the modeled regions; and (3) bilinear parameters encoding the modulations of the specified intrinsic connections by experimental manipulations^62^. The third set of parameters is used to quantify modulatory effects, which reflect increases or decreases in connectivity between two regions given some experimental manipulation, compared with the intrinsic connections between the same regions in the absence of experimental manipulation. fMRI data were modeled using GLM, with regressors for the *Same* condition, and a second condition comprising all visual inputs (i.e., the *Same* and *Different* conditions), which was added specifically for the DCM analysis to be used as the extrinsic visual input.

Given the extrinsic visual input into both TMT+ (the ROI in MT+ evoked by the stimulus on the target side) and IMT+ (the ROI in MT+ evoked by the stimulus on the ignored side), bidirectional connections were hypothesized to exist among the PPC, ACC, IFJ, DLPFC, TMT+, and IMT+. We defined a full model with the modulatory input (the *Same* condition) affecting the bidirectional connections between PPC and ACC, between PPC and IFJ, between PPC and DLPFC, between ACC and IFJ, between ACC and DLPFC, and between IFJ and DLPFC, as well as the intrinsic connection from PPC, from ACC, from IFJ, and from DLPFC to IMT+ (Fig. 5a). We examined this full model for each subject and examined the change in the modulatory effect by the *Same* condition for the post-test, relative to the pre-test (Fig. 5b).

### Eye movement recording

Eye movements were recorded with an EyeLink 1000 Plus system (SR Research, Ltd., Mississauga, Ontario, Canada) in a psychophysics lab (the scanner did not have an applicable eye tracking system). Recording (500 Hz) was performed when subjects performed the same task as psychophysical and fMRI experiments. Supplementary Fig. 2 shows that subjects’ eye movements were small and statistically indistinguishable between the *Same* and *Different* conditions.

## Acknowledgements

We acknowledge the subjects for their contribution to this study. This work was supported by National Outstanding Youth Science Fund Project of National Natural Science Foundation of China (32022032); National Natural Science Foundation of China General Program (31871135 and 32271099); and Key Realm R&D Program of Guangzhou (202007030005).

## Author contributions

X.Z. designed the study. L.H. and J.W., analyzed the data. L.H., J.W., Q.H., C.L., and Y.S. performed research. L.H., J.W., C.S., and X.Z. wrote and edited the paper. All authors evaluated the final manuscript.

