## Supplemental Information for "Category-induced global effects of feature-based attention in human visual system"

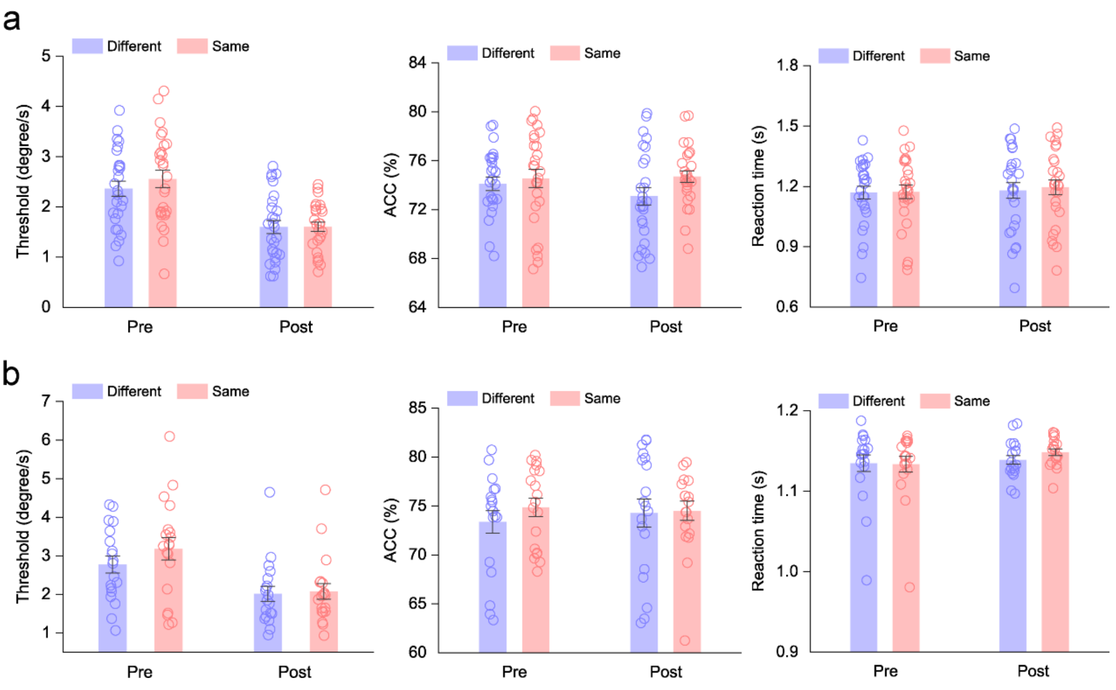


**Fig. S1. Change detection thresholds, response accuracies, and reaction times of the *Same* and *Different* conditions in the pre- and post-tests for psychophysical and fMRI experiments**

**a** In psychophysical experiment, the speed change detection thresholds (mean degree ± SEM) were 2.361 ± 0.149, 2.554 ± 0.172, 1.597 ± 0.129, and 1.602 ± 0.093; the accuracy rates (mean percent correct ± SEM) were 74.108 ± 0.551 %, 74.549 ± 0.740 %, 73.089 ± 0.708 %, and 74.705 ± 0.471 %; and the reaction times (mean reaction time ± SEM) were 1.169 ± 0.031 s, 1.172 ± 0.034 s, 1.180 ± 0.039 s, and 1.195 ± 0.037 s for Pre-Different, Pre-Same, Post-Different, and Post-Same conditions, respectively. All these measurements were submitted to a repeated-measures ANOVA with test (pre and post) and category (*Same* and *Different*) as within-subject factors. Results showed that the main effect of category (thresholds: $F_{1,26} = 0.894$, $p = 0.353$, $\eta_p^2 = 0.033$; accuracy rates: $F_{1,26} = 3.154$, $p = 0.087$, $\eta_p^2 = 0.108$; reaction times: $F_{1,26} = 2.264$, $p = 0.144$, $\eta_p^2 = 0.080$) and the interaction between these two factors (thresholds: $F_{1,26} = 0.844$, $p = 0.367$, $\eta_p^2 = 0.031$; accuracy rates: $F_{1,26} = 2.193$, $p = 0.151$, $\eta_p^2 = 0.078$; reaction times: $F_{1,26} = 1.616$, $p = 0.215$, $\eta_p^2 = 0.059$) were not significant, indicating no significant difference in all these measurements between the *Same* and *Different* conditions. Error bars denote 1 SEM calculated across subjects and colored dots denote the data from each subject. **b** In fMRI experiment, the speed change detection thresholds were 2.774 ± 0.220, 3.185 ± 0.293, 2.016 ± 0.194, and 2.080 ± 0.204; the accuracy rates were 73.363 ± 1.165 %, 74.836 ± 0.945 %, 74.281 ± 1.439 %, and 74.508 ± 0.986 %; and the reaction times were 1.135 ± 0.010 s, 1.133 ± 0.010 s, 1.139 ± 0.005 s, and 1.148 ± 0.004 s for Pre-Different, Pre-Same, Post-Different, and Post-Same conditions, respectively. All these measurements were submitted to a repeated-measures ANOVA with test (pre and post) and category (*Same* and *Different*) as within-subject factors. Results showed that the main effect of category (thresholds: $F_{1,18} = 1.915$, $p = 0.183$, $\eta_p^2 = 0.096$; accuracy rates: $F_{1,18} = 0.450$, $p = 0.511$, $\eta_p^2 = 0.024$; reaction times: $F_{1,18} = 0.564$, $p = 0.462$, $\eta_p^2 = 0.030$) and the interaction between these two factors (thresholds: $F_{1,18} = 2.551$, $p = 0.128$, $\eta_p^2 = 0.124$; accuracy rates: $F_{1,18} = 0.379$, $p = 0.546$, $\eta_p^2 = 0.021$; reaction times: $F_{1,18} = 0.982$, $p = 0.335$, $\eta_p^2 = 0.052$) were not significant, also indicating no significant difference in all these measurements between the *Same* and *Different* conditions. Error bars denote 1 SEM calculated across subjects and colored dots denote the data from each subject.

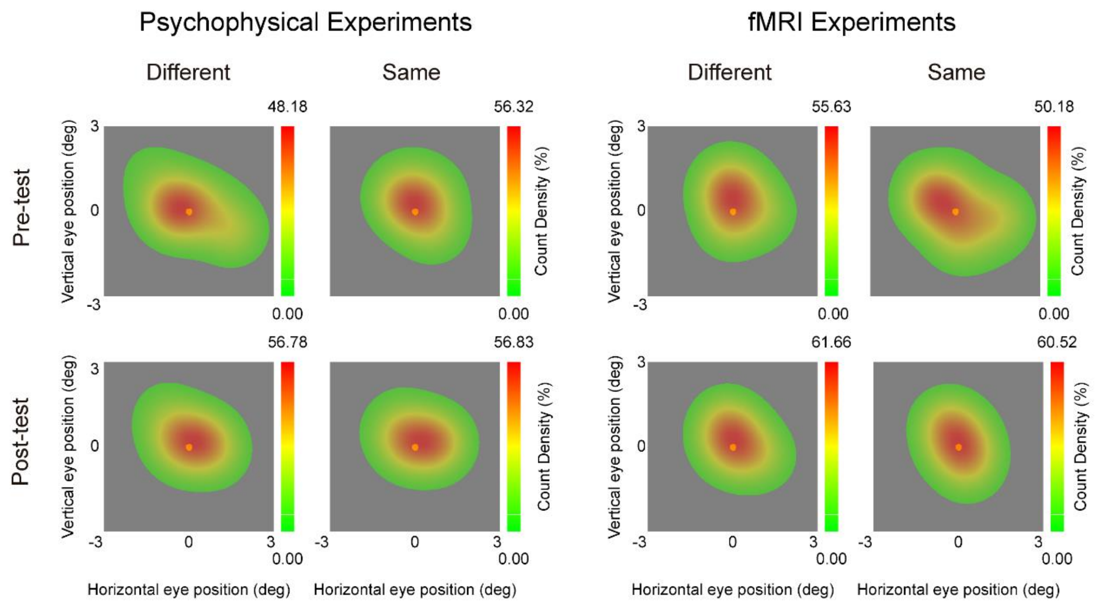


**Fig. S2. Eye movement data in psychophysical and fMRI experiments**

Horizontal and vertical eye positions after removing blinks and artifacts of the *Same* and *Different* conditions in the Pre- and Post-tests for the psychophysical (left) and fMRI (right) experiments. Subjects' eye movements were small (< 3.0°) and not systematically different between the *Same* and *Different* conditions (all $p > 0.05$). Altogether, these results suggest that our results cannot be explained by task difficulty, attention shifting, or eye movement. First, our results cannot be explained by any difference in task difficulty or, presumably, attention, between the *Same* and *Different* conditions. Subjects in our study were asked to detect the speed change of the attended stimulus at threshold, determined by the QUEST staircase procedure (75% correct) before both psychophysical and fMRI tests to ensure that subjects performed equally well for the *Same* and *Different* conditions. Second, our results also cannot be explained by subjects inadvertently shifting their spatial attention to the ignored stimulus in the *Same* condition, as that would have impaired task performance, but there was no significant performance difference between the two conditions. Finally, the eye movement data showed that the subjects' eye movements were small and their eye position distributions were statistically indistinguishable between the *Same* and *Different* conditions. Although our eye movement data were recorded in a psychophysics lab (outside the scanner), it should be noted that the recordings were made when subjects performed the same task as the one in the fMRI experiments. Differences in eye movements for the *Same* and *Different* conditions may be a potential confound, but it is highly unlikely since our recordings outside the scanner did not detect any such differences.
